# Exploring dose-response relationships in *Aedes aegypti* survival upon bacteria and arbovirus infection

**DOI:** 10.1101/2022.09.29.510144

**Authors:** Mariana M. Rocha, Octávio A. C. Talyuli, Clara Luíza Rulff da Costa, Lucilene W. Granella, Daniel S. Mansur, Pedro L. Oliveira, José Henrique M. Oliveira

**Affiliations:** Departamento de Microbiologia, Imunologia e Parasitologia. Universidade Federal de Santa Catarina. Florianópolis, Brazil; Instituto de Bioquímica Médica Leopoldo de Meis. Universidade Federal do Rio de Janeiro. Rio de Janeiro, Brazil; Instituto Nacional de Ciência e Tecnologia em Entomologia Molecular. Brazil

**Keywords:** *Aedes aegypti*, Dengue virus, Zika virus, *Listeria monocytogenes*.

## Abstract

A detailed understanding of host fitness changes upon variation in microbe density is a central aim of infection biology. Here, we applied dose-response curves to study *Aedes aegypti* survival upon exposure to different microbes. We challenged female mosquitoes with *Listeria monocytogenes*, a model bacterial pathogen, Dengue 4 virus and Zika virus, two medically relevant arboviruses, to understand the distribution of mosquito susceptibility and net fitness (survival) following microbe exposure. By correlating microbe loads and host health, we found that a blood meal promotes survival in our systemic bacterial infection model and that mosquitoes orally infected with bacteria had an enhanced defensive capacity than insects infected through injection. We also showed that *Aedes aegypti* has a higher survival profile upon arbovirus infection but, under the conditions tested, was more susceptible to Zika virus when compared to Dengue virus. Here, we applied a framework for the study of microbe-induced mosquito mortality detailing how *Aedes aegypti* lifespan varies upon different inoculum sizes of bacteria and arboviruses.

## 1 Introduction

How microbe density affects host fitness (an ecological measure of disease tolerance) is a central problem in infection biology and microbial pathogenesis (Råberg et al., 2009; Graham et al. 2011). This question has been relatively well-studied in several organisms, from mammals to model insects (Råberg et al., 2007; Louie et al. 2016; Torres et al. 2016; Gupta and Vale 2017; Prakash et al., 2022). In mosquitoes, higher tolerance enhanced vector capacity and transmission of dog heartworm in a natural population of *Aedes albopictus* (Dharmarajan et al., 2019), but the molecular mechanisms of how disease tolerance operates in mosquitoes are only beginning to be explored (Goic et al. 2016). In theory, the manipulation of insect physiology to inhibit disease tolerance could induce vector mortality, providing a biotechnological strategy for arbovirus control (Lambrechts and Saleh 2019; Oliveira et al., 2020).

Here, we applied dose-response curves to study *Aedes aegypti* survival upon exposure to different microbes (bacteria and arbovirus) to understand the distribution of mosquito susceptibility upon infection and net fitness (survival) (Pessoa et al. 2014; Ben-Ami et al., 2010). We infected *Aedes aegypti* through injection (systemic infection) or feeding (midgut infection) with varying doses of the model intracellular pathogen *Listeria monocytogenes* or the epidemiologically relevant flavivirus Dengue and Zika. *Listeria monocytogenes* is an intracellular bacteria able to adapt and grow in various conditions (Toledo-Arana et al. 2009). It is frequently used as a model pathogen (Cossart, 2011) due to its ability to cross anatomical barriers, well-described mechanisms of virulence, and interactions with vertebrate and invertebrate host cells, including insects (Mansfield et al. 2003; Shirasu-Hiza and Schneider 2007). Dengue and Zika are flaviviruses transmitted by mosquitoes and responsible for significant human morbidity and mortality worldwide (Cattarino et al. 2020). How varying initial microbe loads modulate *Aedes aegypti* survival remains to be systematically studied. This question was explored in the following experiments.

## 2 Materials and Methods

### 2.1 Mosquitos

*Aedes aegypti* females (Red Eye strain) were reared and maintained under standard conditions, as described previously (Oliveira et al. 2011; Oliveira et al. 2017). Infections were started in female mosquitoes between 4 and 6 days post adult eclosion.

### 2.2 Bacterial infections

We infected *Aedes aegypti* females with *Listeria monocytogenes* (10403S strain - streptomycin-resistant) (Bécavin et al. 2014). We tested the effect of two different routes: (a) systemic infection through intrathoracic injection or (b) midgut infection through feeding. Frozen bacterial stocks were cultured overnight in brain heart infusion (BHI) media at 37°C without agitation then plated onto LB agar media supplemented with streptomycin (100 ug/mL) and incubated overnight at 37°C. (a) Infection through injection – Before each infection, a single bacterial colony was grown overnight in BHI media at 37°C without agitation until the log phase. Optical densities (OD) at 600 nm were determined and appropriate amounts of culture were centrifuged at room temperature and 1000 x g to pellet bacteria. Bacterial pellets were resuspended in 1mL of phosphate-buffered saline (PBS) to meet the desired ODs (Figure 1 – 0.01; 0.1; 1; 10) and kept on ice until injection. Mosquitoes were cold-anesthetized (for no longer than 20 minutes, including injection time) and injected with 69 nL of bacterial solution in the thorax with a pulled-glass capillary needle attached to Drummond’s Nanoject II microinjector. Infected mosquitoes were transferred to cardboard cups and had *ad libitum* access to cotton pads soaked with 10% sucrose solution (b) Oral infection – A *Listeria monocytogenes* culture was prepared as above and appropriate amounts (Figure 2 – 0.01; 0.1; 1) were pelleted, washed, and resuspended in a protein-rich, chemically-defined substitute blood meal (SBM) that mimics mosquito physiology and digestion when compared to a regular blood meal (Talyuli et al. 2015). Mosquito feeding was performed using water-jacketed artificial feeders maintained at 37°C sealed with parafilm membranes as described elsewhere (Oliveira et al., 2011). Fully engorged females were transferred to cardboard cages with *ad libitum* access to 10% sucrose solution.

**Figure 1:**
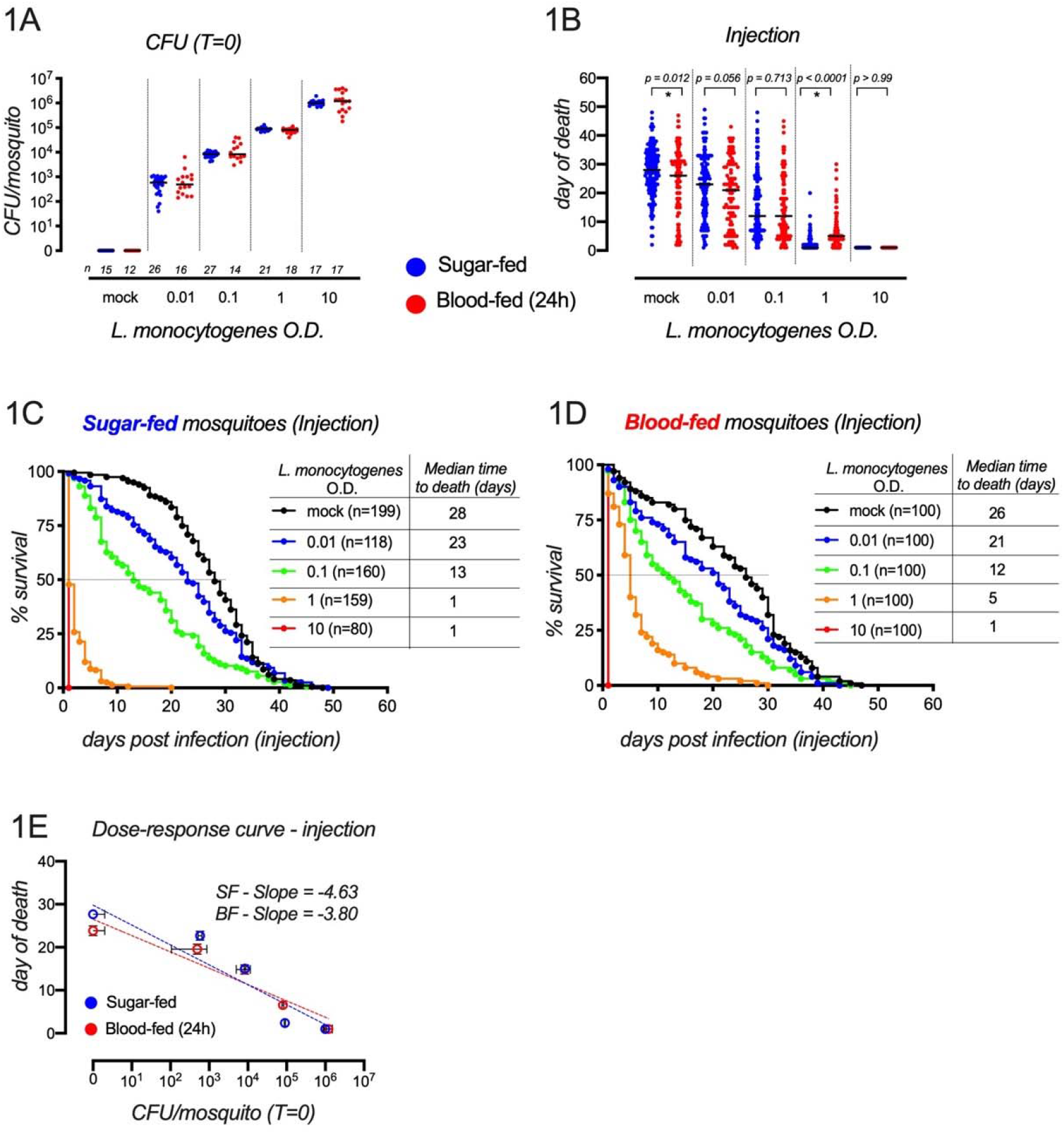
*Aedes aegypti* systemic infection with *Listeria monocytogenes*. (A) CFU analysis: female mosquitoes were injected in the thorax with different amounts of *Listeria monocytogenes* and the number of CFUs determined through spot-plating in LB + Streptomycin agar plates. “Mock” represent PBS-injected mosquitoes. Three independent batch injections were performed and the total number of individual mosquitoes is depicted in the figure. Black bars between individual values represent medians. (B - D) Survival analysis of pooled replicates of at least 3 independent survival curves for every condition tested. Figure B is a summary of data obtained in figures C and D. Black bars between individual values represent medians. In B, *p-values* were determined using the Mann-Whitney test. C – Full survival curve of SF mosquitoes infected with *L. monocytogenes*. Dashed line indicates the median time to death. D – Full survival curve of BF mosquitoes infected with *L. monocytogenes*. (E) Dose-response curves. Data presented in *E* is a correlation between CFU counting (A) and day of death (B). Vertical error bars (survival) and horizontal error bars (CFU) represent the standard error of the mean (SEM). Dashed lines indicate semi-log plots (Graphpad prism V8).

**Figure 2:**
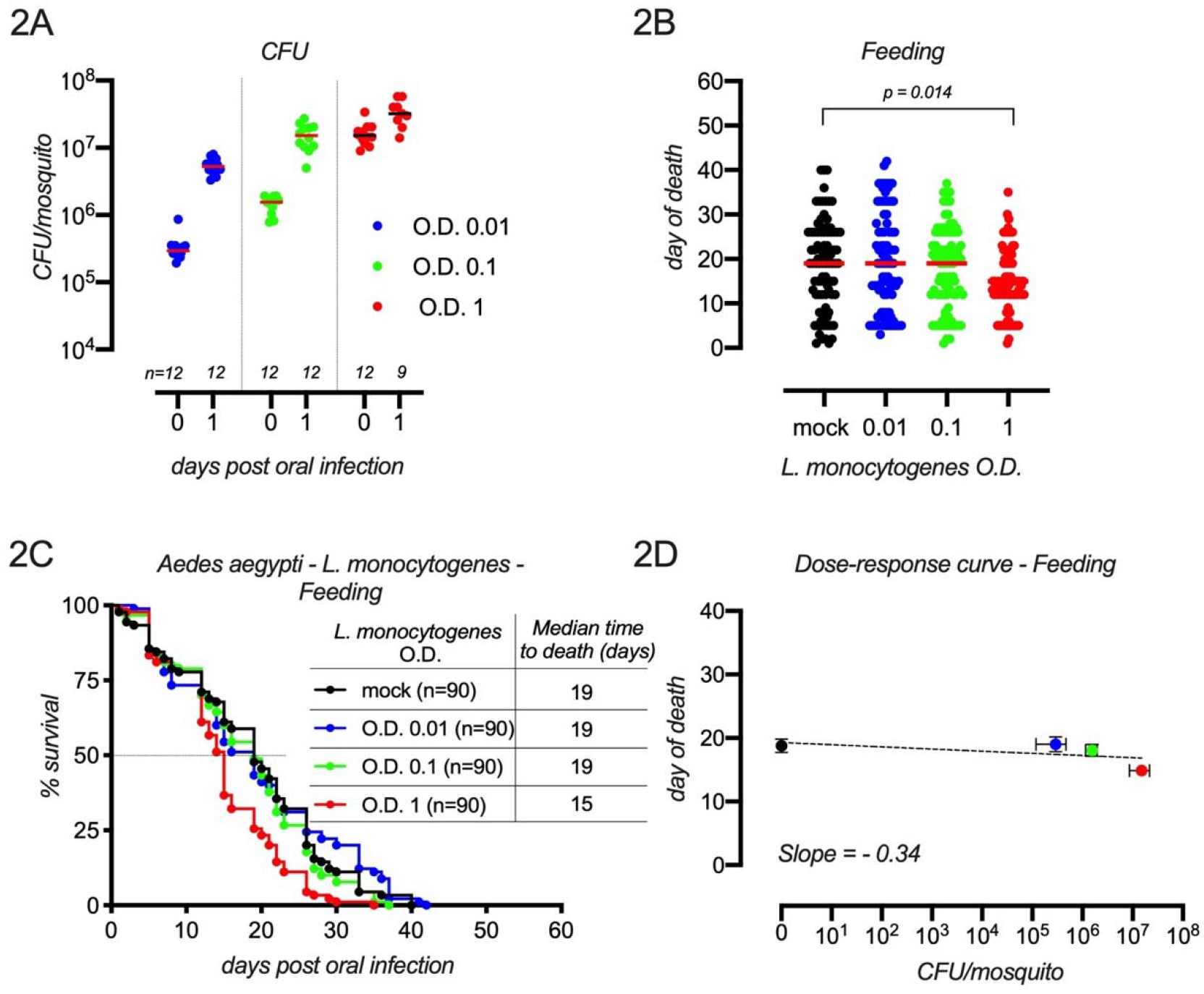
*Aedes aegypti* oral infection with *Listeria monocytogenes*. (A) CFU analysis: female mosquitoes were fed a substitute blood meal (SBM) (Talyuili et al., 2015) supplemented with different amounts of *Listeria monocytogenes* and the number of CFUs determined through spot-plating in LB + Streptomycin agar plates immediately after feeding and 24 hours post-infection. Two independent oral infections were performed for CFU analysis and the number of individual mosquitoes analyzed is depicted in the figure. (B - C) Survival analysis. Figure B is a summary of data obtained in figures C. In B, the *p-value* was determined using the Kruskal-Wallis Multiple Comparison test. Bars between individual values represent medians. (C) Full survival curves of orally infected mosquitoes. Three independent infections were performed and pooled. Dashed line indicates the median time to death. A log-rank (Mantel-Cox) test comparing mock (SBM) vs SBM + *Listeria monocytogenes* O.D. 1 indicated a statistically significant difference in the survival trend between the two groups (p = 0.0002). (D) Dose-response curves. Data presented in *D* is a correlation between CFU counting at T0 (A) and day of death (B). Vertical error bars (survival) and horizontal error bars (CFU) represent the standard error of the mean (SEM). The dashed line indicates a semi-log plot (Graphpad prism V8).

### 2.3 Bacterial loads

For the quantification of *Listeria monocytogenes* after infection, whole mosquitoes were surface-cleaned with 70% ethanol for 1 minute and individually homogenized with a sterile blue pestle and a pestle mixer in 200ul of sterile PBS. Serially diluted homogenates were spot-plated (10ul) in LB agar plates supplemented with streptomycin (100 ug/mL) and incubated overnight at 37°C to allow colony-forming units (CFU) quantification.

### 2.4 Viral infections

We orally infected *Aedes aegypti* with Dengue-4 virus (strain H241 - GenBank: KR011349.2), Dengue-4 virus (strain TVP/360 - GenBank: KU513442.1), and Zika virus (strain PE243/201 - Genbank: KX197192.1). Stock preparation and mosquito infections were performed as detailed in Oliveira et al., 2017. Briefly, females were sucrose-starved overnight and were offered a meal containing a 1:1 mix of rabbit red blood cells and culture supernatants (L-15 media) containing viral particles of infected C6/36 cells. Mosquitoes were allowed to ingest the infectious meal through a parafilm membrane attached to an artificial feeder kept at 37°C for approximately 40 minutes inside a BSL-2 insectary facility. Mosquitoes were then cold-anesthetized and fully engorged females were separated and housed with *ad libitum* access to 10% sucrose-soaked cotton pads.

### 2.5 Viral loads

Immediately following the infectious meal, fully engorged females were frozen at −80°C until use (no more than 3 months). Virus quantification was performed through plaque assays as described in Oliveira et al., 2017. Briefly, whole mosquitoes were individually surface cleaned by soaking the insect for 120 seconds in 70% ethanol + 60 seconds in 1% bleach + 120 seconds in sterile PBS. Under sterile conditions, individual females were homogenized in an Eppendorf with the aid of a blue pestle in 300ul of Dulbecco’s Modified Eagle Medium (DMEM) supplemented with sodium bicarbonate, 1% L-glutamine (200mM), 10% fetal bovine serum, 1% penicillin/streptomycin and 2.5 ug/mL of Amphotericin B. Following homogenization, 700ul of media was added to reach a final concentration of 1 mosquito per mL of media. Eppendorf tubes were centrifuged for 5 minutes at 4°C at maximum speed. Supernatants were serially diluted and plated (100ul – 1/10 of a mosquito) onto BHK-21 cells (for Dengue 4) or Vero cells (for Zika) monolayers in 24-well plates at 70% confluency followed by the addition of semisolid media containing 0.8% methylcellulose (Sigma #M0512. Viscosity 4,000 cP) DMEM containing 2% fetal bovine serum. Plates were incubated at 37°C and 5% COL for 5 days and stained with 1% crystal violet to allow plaque counting. For a detailed description, see Oliveira et al., 2017.

### 2.6 Survival curves

All infections were started in female mosquitoes between 4 and 6 days post adult eclosion. Infected females were cold-anesthetized and placed in cardboard cups (survival cages) at a density of 20 females per cup (470 mL maximum capacity). 10% sucrose was provided *ad libitum* through cotton pads placed on the top of a woven mesh and replaced 2-3 times a week. Survival was recorded once per day 6 times a week until all the insects inside the cups were dead. Survival cages were kept at the insectary at 28°C (+/− 10%) and 80% humidity (+/−10%). All survival curves are presented as pooled data from at least 3 independent experiments.

### 2.7 Statistical analysis

All survival experiments were performed at least 3 independent times. Replicate numbers are detailed in figure legends. For survival curves, independent replicates were pooled an analyzed using the Logrank test for trend and Gehan-Breslow-Wilcoxon test (GraphPad Prism 8). When appropriate, “day of death” comparisons were performed with Mann-Whitney test or the Kruskal-Wallis Multiple Comparison test, as indicated in figure legends.

## 3 Results

### Listeria monocytogenes as a model pathogen to study Aedes aegypti immunity

We took advantage of the natural resistance to streptomycin of the *Listeria monocytogenes* strain 10403S (Bécavin et al., 2014) to perform dose-response curves and test mosquito survival upon systemic infection of a wide spectrum of initial doses (Figure 1). We observed that a linear increase in the concentration of injected bacteria led to a linear and proportional ability to quantify *Listeria monocytogenes* CFU in whole-body mosquitoes immediately upon injection (Figure 1A). This was true for sugar-fed (SF) and blood-fed (BF) mosquitoes that were offered a blood meal 24 hours before. Next, we measured the net fitness consequence (survival) of *Aedes aegypti* exposed to varying doses of systemically-injected *Listeria monocytogenes* (Figures 1B, 1C, 1D). We observed a wide spectrum of pathogenicity, with a dose-dependent reduction in the median time to death (MTD) and a comparable response between SF and BF insects. The largest difference in MTD of SF and BF mosquitoes was observed in insects injected with OD 1 (SF – MTD = 1 day vs BF - MTD = 5 days). These two groups had a highly significant difference in the distribution of mortality, plotted here as the day of death (Figure 1B – OD 1 - p < 0.0001). Of note, the day-of-death distribution was also statistically different between uninfected mosquitoes (Figure 1 – mock - p = 0.012) suggesting that the different physiological status of SF and BF insects (Sterkel et al. 2017) affected the lifespan of mock-injected mosquitoes (SF – MTD = 28 days vs BF – MTD = 26 days). We followed bacterial load overtime in SF females injected with intermediary doses (OD 0.01 and 0.1) of *Listeria monocytogenes* (Supplementary Figure 1) and observed a high level of variation in infection intensity (number of CFUs per mosquito) and prevalence (proportion of infected mosquitoes regardless of CFU amount), as was previously reported in experiments tracking within-host dynamics of bacterial load in infected insects (Duneau et al. 2017; Graham and Tate 2017; Duneau and Ferdy 2022). Next, we correlated microbe loads immediately following injection (Figure 1A) with the median time to death of SF and BF mosquitoes (Figures 1B-D) to observe how differences in microbe exposure impacted survival. We assumed a linear relationship between microbe load and host survival and fit data using a semilog plot (Simms, 2000). We observed slightly different negative slopes between SF and BF mosquitoes (SF = −4.63 vs BF = −3.80) suggesting that a blood meal increases mosquito survival over a range of *Listeria monocytogenes* initial infection load (Figure 1E). BF insects had reduced vigor (health of non-infected mosquitoes) (Louie et al., 2016; Oliveira et al, 2020) and enhanced survival ability at ~10□ injected CFUs (Figure 1E). Different than BF mosquitoes, SF mosquitoes were equally susceptible at 10□ and 10□ injected CFUs, suggesting possible non-linear relationships between microbe load and host mortality (Doeschl-Wilson et al., 2012; Louie et al. 2016; Gupta and Vale 2017).

Because *Listeria monocytogenes* is mainly a foodborne pathogen and *Aedes aegypti* gets infected frequently by feeding (ex: by arboviruses), we decided to evaluate the effect of varying infectious doses through an oral infection model. We offered different amounts of *L. monocytogenes* mixed with a protein-rich diet, called substitute blood meal (SBM), that mimics mosquito intestinal physiology and digestion when compared to a regular blood meal (Talyuli et al. 2015). We measured bacterial load in whole mosquitoes at two points; immediately after oral feeding (T0) and 24 hours after infection (T24). Similar to injection, feeding different bacterial amounts led to a proportional increase in bacterial load at T0 (Figure 2A). At T24, all tested mosquitoes were infected and allowed bacterial proliferation, ranging from 18X growth between T0 and T24 at OD 0.01, 9X at OD 0.1, but only 2X at OD 1, probably approaching the carrying capacity of *Aedes aegypti* for *Listeria monocytogenes* (Figure 2A). Infection through feeding resulted in a less virulent outcome and a reduced capacity to induce mosquito mortality, with only the highest dose significantly, but modestly, decreasing the median time to death (Figure 2B, 2C). A dose-response curve comparing infection intensity and survival presented an almost flat reaction norm (slope = −0.34 - semilog plot), suggesting *Aedes aegypti* have a higher defense capacity against *L. monocytogenes* when challenged through feeding (Figure 2D) when compared to a systemic infection (Figure 1E).

### Aedes aegypti survival following an arbovirus-infected blood meal

We fed *Aedes aegypti* females with blood supplemented with different amounts of two laboratory-adapted reference strains of Dengue 4 virus, strain H241 (Oliveira et al. 2017), strain TVP/360 (Kuczera et al. 2016), and Zika virus (ZIKV) (Figure 3) and measured viral load at T0 (Figure 3A, 3D, 3G) and mosquito mortality. Under the experimental conditions tested, none of the DENV concentrations offered to mosquitoes resulted in significant lifespan reduction (Figures 3B, 3C, 3E, 3F – Log-rank (Mantel-Cox) test). When fed different concentrations of ZIKV, our experiments revealed a statistically significant reduction in MTD in the group fed the highest dose (10□ PFU/mL of blood) (mock – MTD = 30 days vs 10□ PFU/mL – MTD = 24 days – Log-rank (Mantel-Cox) test) (Figures 3H and 3L). Interestingly, mosquitoes challenged with intermediary doses (10^3^ PFU/mL and 10□ PFU/mL) also displayed minor reductions in lifespan compared to mock (mock – MTD = 30 days vs 10^3^ PFU/mL – MTD = 28 days vs 10□ PFU/mL – MTD = 28 days).

**Figure 3:**
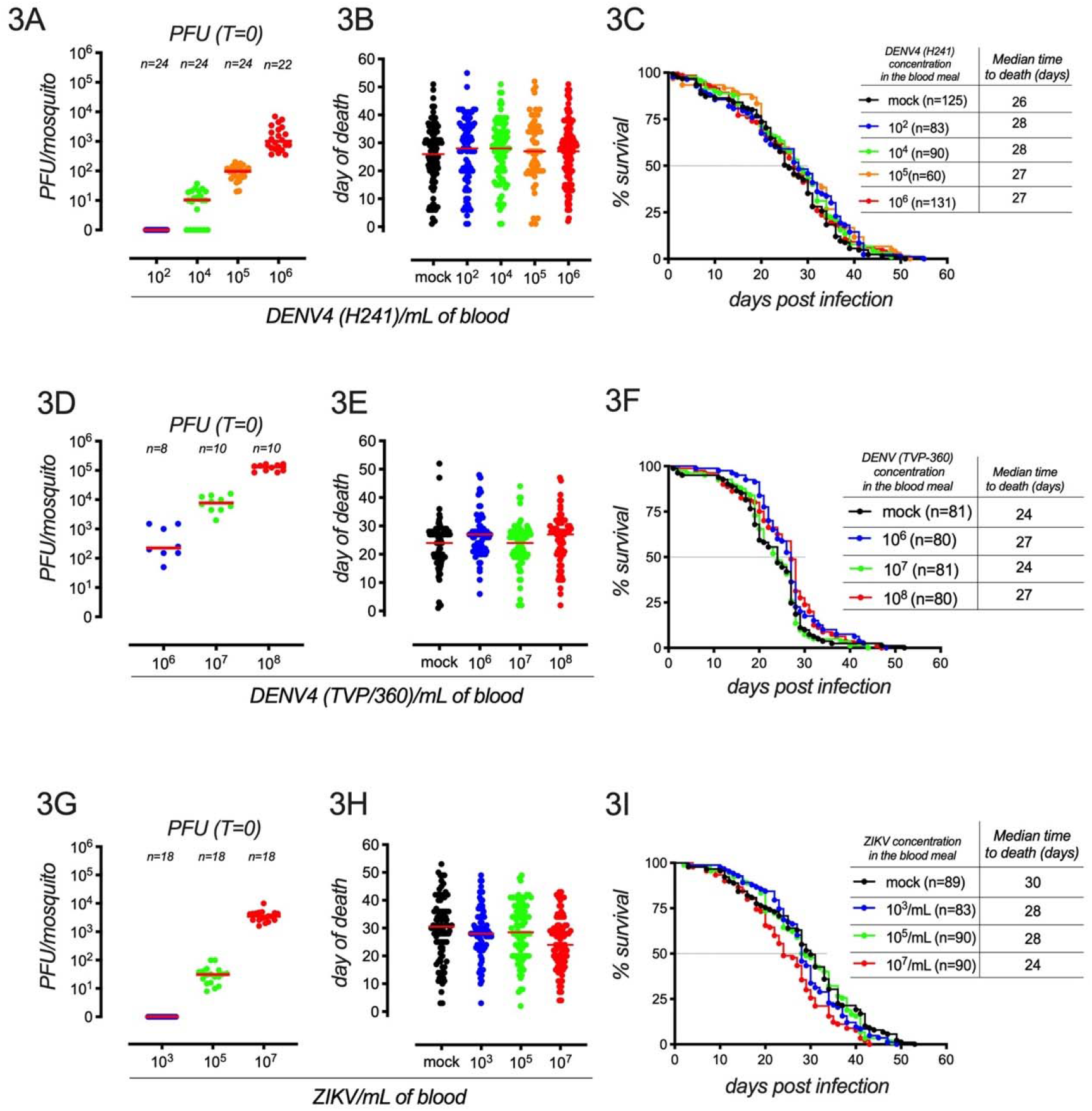
*Aedes aegypti* infected with arboviruses. (A, D, G) Plaque forming unit (PFU) analysis. Virus load was determined immediately after feeding in fully engorged whole-body mosquitoes following 3 independent infections. Bars between individual values represent medians. (B, C, E, F, H, L) Survival analysis of pooled mosquitoes from 3 independent infections. Bars between individual values represent medians (3B, 3E, 3H). 3C, 3F, 3L represent the full survival curve of virus-challenged mosquitoes. Dashed lines indicate the median time to death. Log-rank (Mantel-Cox) tests were used for survival analysis. “Mock” represent a group of mosquitos fed with a 1:1 mix of rabbit red blood cells and culture supernatants (L-15 media) of uninfected C6/36 cells.

Our dose-response analysis demonstrated shallow slopes for both strains of DENV4 (H241: slope = 0.23 – semilog plot and TVP/360: slope = 0.32 - semilog plot) compared to ZIKV (slope = −1.25 - semilog plot), suggesting that, under the conditions tested, DENV4 exposure had a reduced impact on *Aedes aegypti* survival when compared to ZIKV (Figure 4).

**Figure 4:**
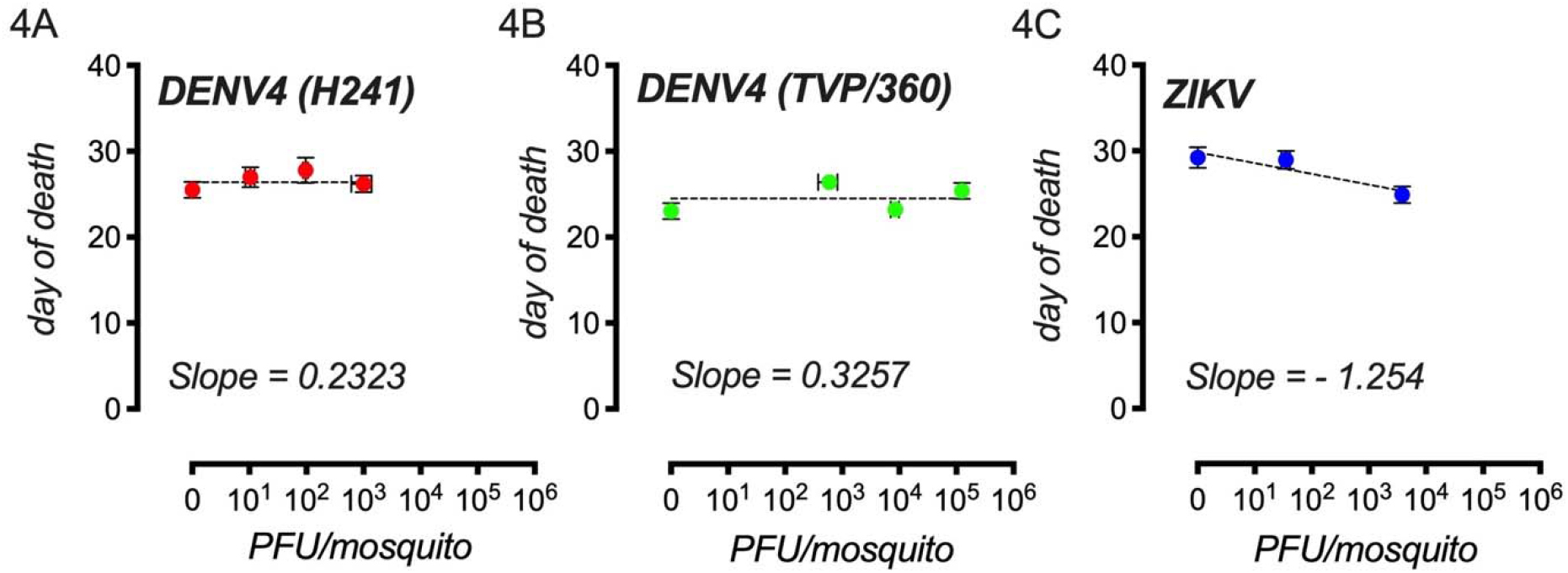
Dose-response curves of arbovirus-challenged mosquitoes. The data presented is a correlation between PFU counting (Figure 3A, 3D, 3G) and the day of death (Figure 3B, 3E, 3H). Vertical error bars (survival) and horizontal error bars (PFU) represent the standard error of the mean (SEM). Dashed lines indicate semi-log plots (Graphpad prism V8).

## Discussion

We applied an eco-immunology approach to investigate *Aedes aegypti* susceptibility to bacteria and arbovirus challenges. By correlating microbe loads and host health (survival) (Simms, 2000), we identified different susceptibility profiles of *Aedes aegypti* during infection (Figure 1E, Figure 2D and Figure 4). This is a novel framework to study mosquito pathogenesis, resistance and disease tolerance (Oliveira et al., 2020; Magistrado et al., 2023). Disease tolerance has been poorly explored in mosquitoes and, as a defensive strategy with the potential to modulate insect fitness, it is likely to influence the selection and evolution of immune traits relevant to vector competence (Schmid-Hempel 2005; Seal et al., 2021). Our results provide a simple descriptive experimental design for the study of infection biology in *Aedes aegypti*. This approach may be useful in mechanistic studies aiming to uncover genes and pathways involved in vector disease tolerance and resistance (Lambrechts and Saleh 2019; Oliveira et al., 2020).

*Listeria monocytogenes* is a model pathogen frequently used in *Drosophila melanogaster* infection studies, but only rarely used to perturb mosquito immunity (Kim et al. 2020). By triggering a systemic infection through injection (Figure 1), we showed different susceptibility profiles of SF and BF mosquitoes (Figure 1E). When comparing infection routes, our data demonstrated enhanced survival upon midgut infection of *L. monocytogenes* (Figure 1E vs 2D). The mechanistic basis of higher survival in BF mosquitoes as well as midgut infections remains to be determined. We speculate that the microbiota, the peritrophic matrix, and stress response pathways might enhance the mosquito defensive capacity in the midgut of blood-fed *Aedes aegypti* (Talyuli et al. 2022; Hixson et al. 2022).

In our dose-response susceptibility curves upon arbovirus challenge, we observed that, under the conditions tested, *Aedes aegypti* exhibited higher pathogenesis when fed with ZIKV, compared to DENV4 (Figure 4). These phenotypes deserve further investigation since, in a different study, DENV2 reduced mosquito survival under similar experimental settings (Maciel-de-Freitas et al., 2011). Distinct mosquito strains, virus serotypes, and virus titers, among other causes, might explain the observed differences. For ZIKV challenges, we observed a 4-day reduction in the median time to death compared to mock-infected (Figure 4C). Since minor changes in vector mortality rate might account for larger differences in the basic reproduction number (RL) (Luz et al. 2003), the observed 4-day reduction in MTD (~15% lifespan) might be relevant in the context of arbovirus epidemics.

One limitation of our results is that assessments of microbe density were measured immediately after exposure. We choose this time point because founder effects are known to determine disease outcomes and vector competence (Duneau et al. 2017; Hodoameda et al., 2022) and our main goal was to assess *Aedes aegypti* survival distribution upon bacteria and arbovirus challenges. Further experiments will employ the dose-response framework used here to quantify vector tolerance against different pathogens and physiological conditions.

## Supporting information

Figura S1

## Acknowledgment

We thank Pedro F. Vale (University of Edinburgh) and André Báfica (Universidade Federal de Santa Catarina) for helpful discussions.

---

**Supplementary figure 1.**
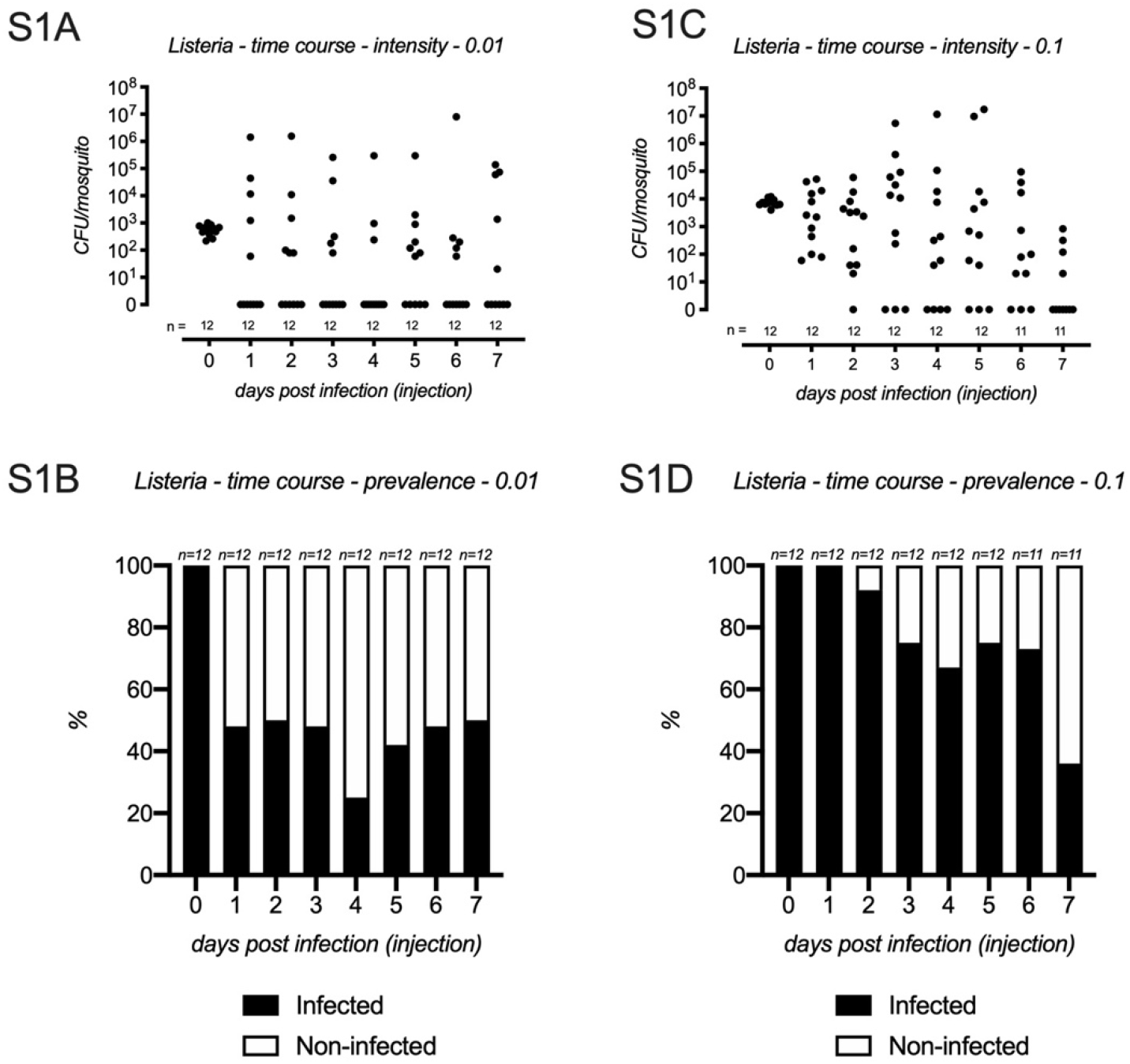
Time-series analysis of *Listeria monocytogenes* infection load in *Aedes aegypti* (SF). Mosquitoes were infected through injection with different amounts of bacteria and monitored over time for CFU counting in whole mosquitoes. Infection intensity (S1A and S1C) is defined as the number of CFU per mosquito. Infection prevalence (S1B and S1D) is defined as the number of *Listeria monocytogenes*-positive mosquitoes, regardless of CFU counts. The number of mosquito replicates is depicted in the figure and is a result of two independent batch infections.

