## Supplementary figures and images for "Exploring dose-response relationships in *Aedes aegypti* survival upon bacteria and arbovirus infection"

### Figura S1

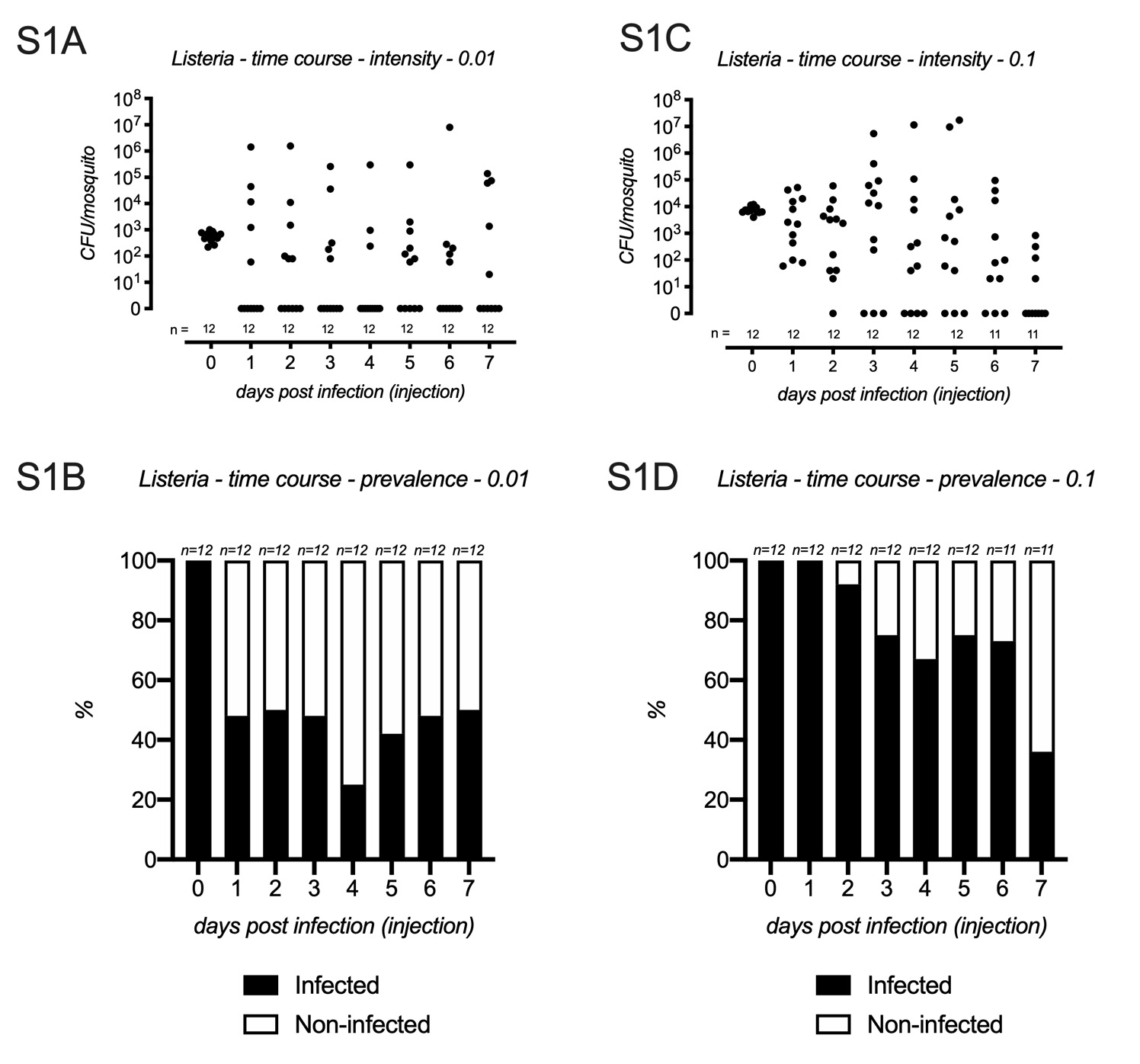
